# Proteomic analysis of *Caenorhabditis elegans* against *S.* Typhi toxic proteins

**DOI:** 10.1101/845370

**Authors:** Dilawar Ahmad Mir, Krishnaswamy Balamurugan

## Abstract

**Background & Aims:** Bacterial effector molecules are the crucial infectious agents and are sufficient to cause pathogenesis. In the present study, pathogenesis of *S.* Typhi toxic proteins on the model host *Caenorhabditis elegans* was investigated by exploring the host regulatory proteins during infection through quantitative proteomics approach.

**Methods:** In this regard, the host proteome was analysed using two-dimensional gel electrophoresis (2D-GE) and differentially regulated proteins were identified using MALDI TOF/TOF/MS analysis. Out of the 150 regulated proteins identified, 95 proteins were appeared to be downregulated while 55 were upregulated. Interaction network for regulated proteins was predicted using STRING tool.

**Results:** Most of the downregulated proteins were found to be involved in muscle contraction, locomotion, energy hydrolysis, lipid synthesis, serine/threonine kinase activity, oxidoreductase activity and protein unfolding and upregulated proteins were found to be involved in oxidative stress pathways. Hence, cellular stress generated by *S.* Typhi proteome on the model host was determined using lipid peroxidation, oxidant and antioxidant assays. In addition to that the candidate proteins resulted from the host proteome analysis were validated by Western blotting and roles of several crucial molecular players were analyzed *in vivo* using wild type and mutant *C. elegans*.

**Conclusions:** To the best of our knowledge, this is the first study to report the protein regulation in host *C. elegans* during *S.* Typhi toxic proteins exposure which highlights the significance of p38 MAPK and JNK immune pathways. These results may provide new clues for future therapeutic approaches for curing bacterial toxin protein-mediated infections in a host system.

**Summary:** We have precipitated the toxin proteins of *S.*Typhi. To gain insight into the worm’s response to ingestion of toxin, a proteomic analysis was performed to monitor the changes in protein regulation. 150 differential regulated proteins were identified, amongst 95 and 55 proteins were found to be downregulated and upregulated, respectively. This is the first study that reported the global proteome changes in *C. elegans* against toxin. Our findings suggested involvement of several regulatory proteins that appear to play a role in various molecular functions in combating toxin-mediated microbial pathogenicity and/or host *C. elegans* immunity modulation. A proteomics approach using *C. elegans* can facilitate the understanding of how toxin can lead to intoxication, which pave a way for delineating how higher eukaryotes could evolve defenses to protect against bacterial toxin. Toxin infection nematode showed increased accumulation of proteins that respond to oxidative –stress, lipid metabolism, embryonic development, immune and inflammatory processes.

## 1. Introduction

*C. elegans* is a well suited model to investigate the cellular impact of bacterial toxic proteins since the model has been well established for its utility in toxicological studies. Its short generation time, large brood size, conventional biology and well defined innate immune system have made it as a suitable model for toxicological studies (Popham *et al*., 1979; Hodgkin *et al*., 2000; Leung *et al*., 2008). Toxicity analysis on *C. elegans* provides the data of whole animal with intact and metabolically active reproductive, digestive, endocrine, sensory and neuromuscular systems (Hunt *et al*., 2017). Most importantly, *C. elegans* provides a naive and rapid biological system to investigate bacterial toxins and host interaction as it naturally feeds on bacteria (Huffman *et al*., 2004). Many Gram-negative and Gram-positive bacteria which are pathogenic to humans are also reported to infect *C. elegans* (Aballay *et al*., 2003). Bacterial pathogens promote host pathogenesis by prominent virulent factors including lipopolysaccharide, flagella, pili, proteases, exotoxin A and exoenzymes (Lyczak *et al*., 2000). Phenazine (virulence factors) toxicity through oxidative stress was first identified in *C. elegans* (Cezairliyan *et al*., 2013) which was later reported in Drosophila, mice and plants also (Ray *et al*., 2015). The viable cultures of *S.* Typhi, *P. aeruginosa* and *K. pneumonia* and their isolated lipopolysaccharide (LPS) act as a powerful immune activator and lethal agent on this nematode model (Vigneshkumar *et al*., 2012; Sivamaruthi *et al*., 2014; Kamaladevi *et al*., 2016). It was reported that the lipoteichoic acid of Gram-positive bacteria is equivalently antigenic to LPS which elicit the inflammatory response in the host (JebaMercy *et al*., 2015).

Our previously reports attest the utility of *C. elegans* as a model organism for the following human pathogens, *Shigella* spp, *Vibrio alginolyticus, Proteus spp.* and *S.* Typhi (Kesika *et al*., 2011; Durai *et al*., 2011; Kesika and Balamurugan 2012; Jebamercy *et al*., 2013; Sivamaruthi *et al*., 2014). These reports uncover the molecular players responsible for host defence upon *K. pneumonia* and *P. aeruginosa* pathogenesis through proteomic approach (Balasubramanian *et al*., 2016; Kamaladevi and Balamurugan 2017). *Salmonella* spp., are responsible for millions of infections per year ranging from food poisoning to life-threatening systemic typhoid fever (Gal-Mor *et al*., 2014). For pathogenesis, *Salmonella* spp. uses Type-III Secretion System (T3SS), to deliver bacterial proteins/toxins directly into eukaryotic host cells and amends its cellular functions (Galan *et al*., 2001). In particular, *S.* Typhi genome encodes ∼ 4500 proteins identified by proteome analysis (Liu *et al*., 2015). Toxic proteins are poisonous substance and are capable of causing diseases on contact with or absorption by host body tissues, these proteins interact with biological macromolecules such as enzymes or cellular receptors and disable the host immune system (Smith *et al*., 1972; Schlesinger D 1975). Hence, in the current study we have investigated the impact of whole proteome of *S.* Typhi on translational machinery of *C. elegans* by employing proteomic approaches. In this context, *S.* Typhi whole-cell enriched proteins were used to treat *C. elegans* and subsequently the host proteome was isolated and analysed using 2D-GE. To the best of our knowledge, this is the first study to identify differentially regulated candidate proteins in *C. elegans* against bacterial toxins using proteomic approach.

## 2. Results

### 2.1 Intact protein of *S.* Typhi is essential to elucidate pathogenesis in *C. elegans*

The *S.* Typhi and *E. coli* OP50 proteome were precipitated using (NH4)_2_SO_4_ and confirmed by resolving in SDS-PAGE. The protein bands with varying intensity and pattern were seen in precipitated fractions (**Supplemental Figure 1**). To determine the effect and importance of bacterial proteins on the host-pathogen interaction, killing ability of *S.* Typhi and *E. coli* OP50 precipitated protein was assessed by performing killing assay. The result indicated that *S.* Typhi proteome required 36 ± 5 hrs (*p* < 0.05) for the complete killing of *C. elegans* (**Figure 1A**) with the LT_50_, (time for half to die) of 20 ± 2 hrs. Ammonium sulphate precipitated protein fractions (50% – 80%) killed the *C. elegans* with mean life span 36 ± 5 hrs, whereas *E. coli* OP50 proteome fraction (50% – 70%) at same concentration (1.5 mg/mL) neither killed the worms nor was able to bring any significant physiological changes on a relevant time scale. To confirm whether the toxin proteins or any other bacterial agent are responsible for the *C. elegans* mortality, precipitated protein fractions of *S.* Typhi was digested with proteinase-K (broad-spectrum of serine proteases which are able to digest the proteome) overnight at 37°C and tested for its pathogenicity. The result clearly denoted that there was no significant (*p* < 0.05) difference between mean life span in overnight digested protein fractions and control worms fed with *E. coli* OP50 which suggested that only the protein fractions have modulated the *C. elegans* lifespan and morphology **(Figure 1B**). The lethal *S.* Typhi toxin protein fractions (50% – 80%) at 1.5 mg/mL, 1 mg/mL and 500 μg/mL concentration significantly (*p* < 0.05) killed *C. elegans* at 36, 65, and 90 hrs respectively, whereas *E. coli* OP50 and its proteome fraction has not bring any significant physiological changes in the nematode (**Figure 1C**). To confirm the (NH4)_2_SO_4_ effect on nematode, the L4 stage N2 worms were grown in various concentrations of (NH4)_2_SO_4_ medium and monitored for their survival rate. It was found that (NH4)_2_SO_4_ medium exceeding the concentration of 500 mM are lethal to *C. elegans* (**Figure 1D**). Each experiment was performed in biological triplicates and the error bars represent the mean ± SD (\**p* < 0.05).

**Figure 1.**
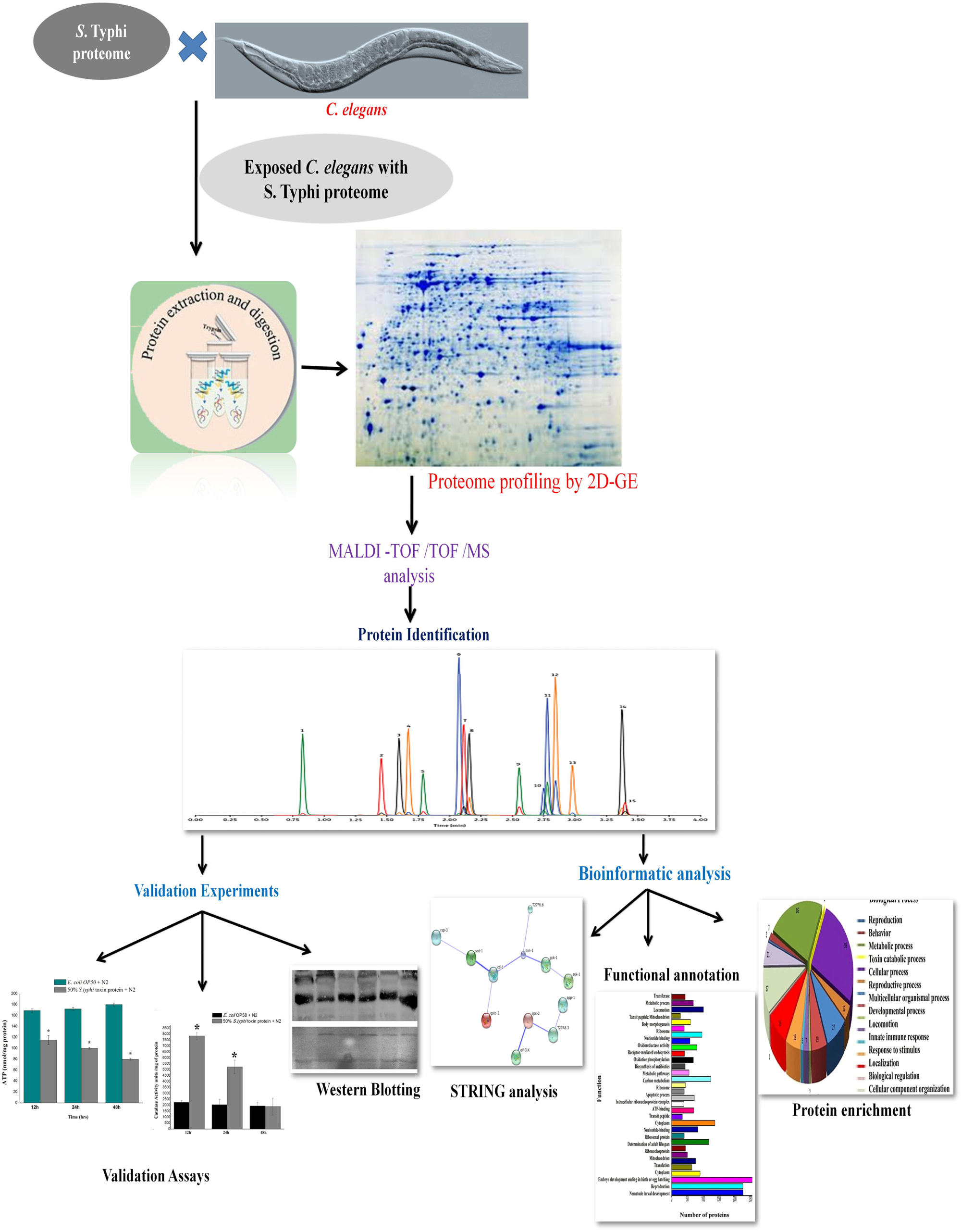
**A**) Physiological assays showing the impact of *E. coli* OP50 and *S.* Typhi toxin proteins on wild type *C. elegans*. In liquid killing assays, 50% – 80% Ammonium Sulphate (AS) precipitated *S.* Typhi toxin proteins caused complete killing of *C. elegans* at 36 ± 4 hrs. **B)** Toxin protein fractions (50% – 80%) digested by Proteinase-K has not killed the nematode. Whereas Proteinase-K untreated protein fraction (50% – 80%) killed the host and confirmed that the proteins are only responsible for the *C. elegans* mortality. **C**) The *S.* Typhi toxin protein at 1.5 mg/mL, 1.0 mg/mL and 500 μg/mL concentrations killed *C. elegans* significantly (*p* < 0.05) at 36, 65, and 90 hrs respectively, whereas *C. elegans to E. coli* OP50 total protein showed normal life span. **D**) Medium containing AS concentration (200 – 800 mM/mL) was used to check its toxicity against nematode. Medium exceeding the AS concentration at 500 mM/mL is lethal to *C. elegans*. Each experiment was performed in triplicates. *Differences were considered significant at *p* < 0.05.

### 2.2 2D-GE based proteomic analyses of *C. elegans* upon exposure to toxins

The results of the killing assay and protein pattern of *C. elegans* on SDS-PAGE (**Supplemental Figure 2**) have lead us to investigate the primary molecular mechanism of *C. elegans* mortality through 2D-GE. Worms exposed to *S.* Typhi toxins for 24 hrs were taken for the analysis. A 2D-GE was deployed to decipher protein regulation in control and treated samples respectively (**Figure 2A and B**). The 2D-GE triplicate gels are shown in Supplemental Figure 3. The protein spots present in the control and treated 2D-GE gels were matched and compared using Image Master Platinum 7 software (GE Healthcare). Based on the densitometry analysis 477 detected protein spots were found to satisfy the arbitrary parameters. The relative expression ratio of downregulated and upregulated protein spots in control and treated sample was fixed at ≥ **-**1.5 and ≥ 1.5 respectively of all the biological replicates (*p* < 0.05). Among 477 matched spots, 95 and 55 spots were found to be downregulated and upregulated, respectively. Selected differentially expressed protein spots were excised from preparative gel and analysed by MALDI-TOF/TOF/MS, and proteins were identified by Mascot tool. The list of identified differentially regulated proteins with their Mascot score, percentage of sequence matched and fold change is provided in **Supplemental Tables 1 and 2**. A GO classification of regulated proteins was performed using the UniProtKB tool to categorize regulated proteins into, biological processes, molecular functions and cellular components. The functional annotations of largest set of regulated proteins are presented in (**Figure 3**). Most of the identified regulated proteins have shown to play important roles in embryology, cytoskeleton, reproduction, metabolism development, ubiquitination, and oxidative stress. All these biological processes are directly reliable response to changes made by stress conditions. The interaction among regulated protein players of *C. elegans* was performed using the STRING tool. The interaction analysis was performed to decipher high degree of connectivity and their role in important biological pathways. The interaction map displayed the relation between the identified regulated proteins of *C. elegans* as presented in (**Figure 4A**). The downregulated protein DAF-21 showed interaction with MYO-1/2, UNC-54, LET-754, H28O16.1, STI-1, KGB-1, TRX-1 and HSP-16.2 proteins. These proteins play an important role in muscle contraction (*myo-1/2*), locomotion (*unc-54*), energy hydrolysis (*let-754*), ATP synthase complex (H28O16.1), stress response (*sti-1*), serine/threonine kinase activity (*kgb-1*), oxidoreductase activity (*trx-1*) and protein unfolding (*hsp-16.2*). STRING analysis of upregulated proteins displayed the interaction between SOD-1, CTL-1, PXN-1, GCK-1 and SEK-1 which is ubiquitously related to oxidative stress. Functional annotation and gene enrichment of all regulated proteins [N=150] were performed using DAVID tool. Functional annotation reflects the character of one protein in diverse biological processes. The biological functions whose highest numbers of proteins that are differentially regulated in response to *S.* Typhi proteome exposure were embryonic development [N=56], post embryonic development [N=38] as provided in (**Figure 4B**). Biological process that exhibits highest gene enrichment score were cytoskeleton, metal binding, cell adhesion, protease and redox processes as provided in (**Figure 4C**).

**Table 1.** *In vivo* embryo development assay. From gravid *C. elegans*, worm embryos were isolated treated to *S.* Typhi protein fraction (concentration 1.0 – 1.5 mg/mL) and *E. coli* OP50 (control). Treated embryos has not transformed into L1 larval developmental stage of *C. elegans* up to 48 hrs whereas embryo treated with *E. coli* OP50 (control) grows normally after 12 hrs of incubation (**Table 1**).The positive and negative signs given in a table shows presence or absence of L1 larval stage. This *in vitro* embryo development assay was performed in triplicates.

| Serial No. | 12h L1<br>Development | 24h L2<br>Development | 48h L4<br>Development |
| --- | --- | --- | --- |
| *E. Coli* OP50 bacteria<br>(Control) | + | + | + |
| 50% *S. Typhi* toxin protein<br>(1.5mg/mL concentration) | - | - | - |
| 50% *S. Typhi* toxin protein<br>(1.0mg/mL concentration) | - | - | - |

+ = Present
- = Absent
L1= First Larval Stage of *C. elegans*

**Figure 2.**
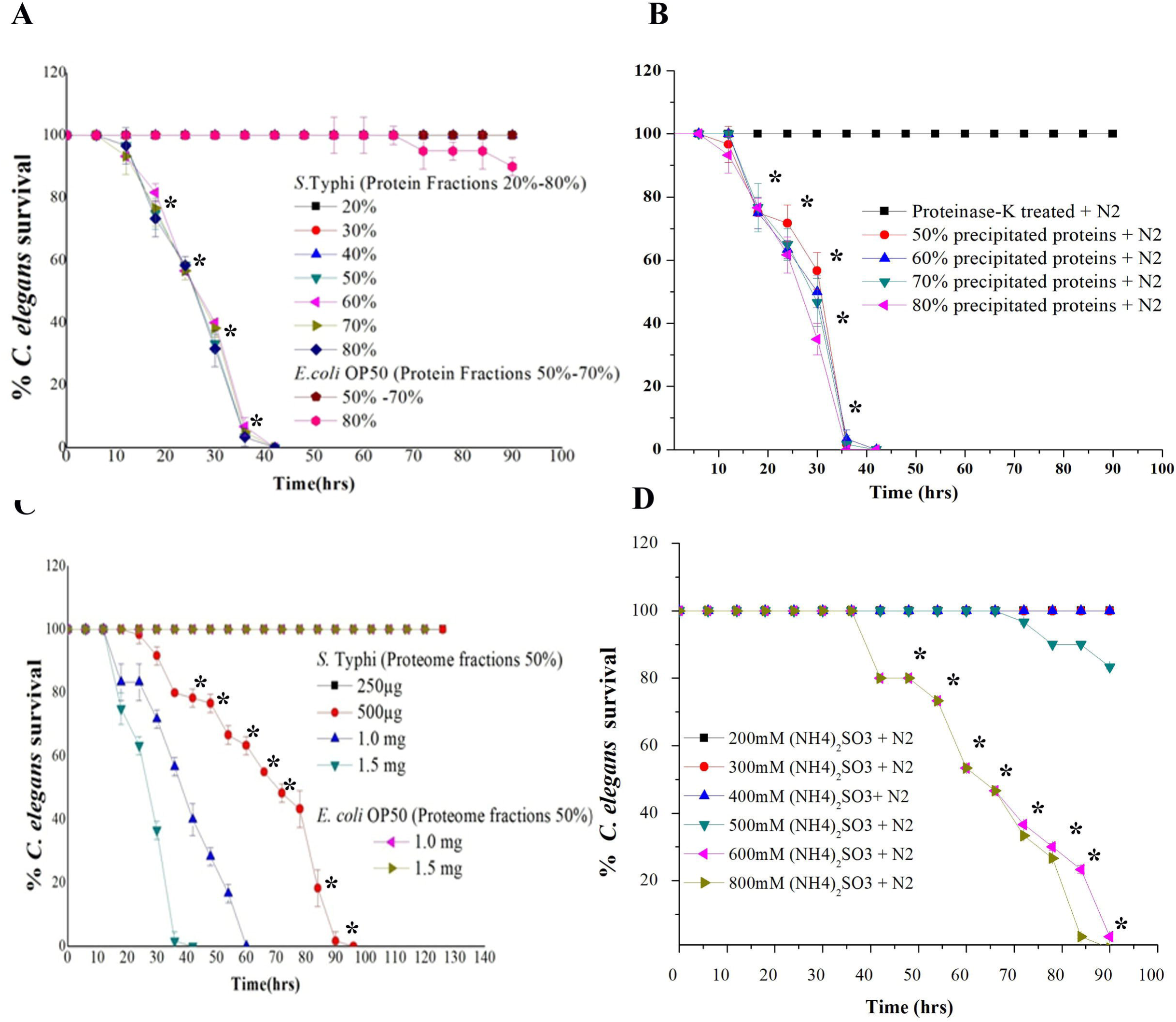
2D gel electrophoreses images of *C. elegans* proteome. **A**) The control nematodes fed with *E. coli* OP50. **B**) *C. elegans* total proteins treated to 1.5 mg/mL of 50% *S.* Typhi protein fraction. The size of IP strip is 18 cm and the pI gradient is from 3 – 10. The experiment was performed in triplicates.

**Figure 3.**
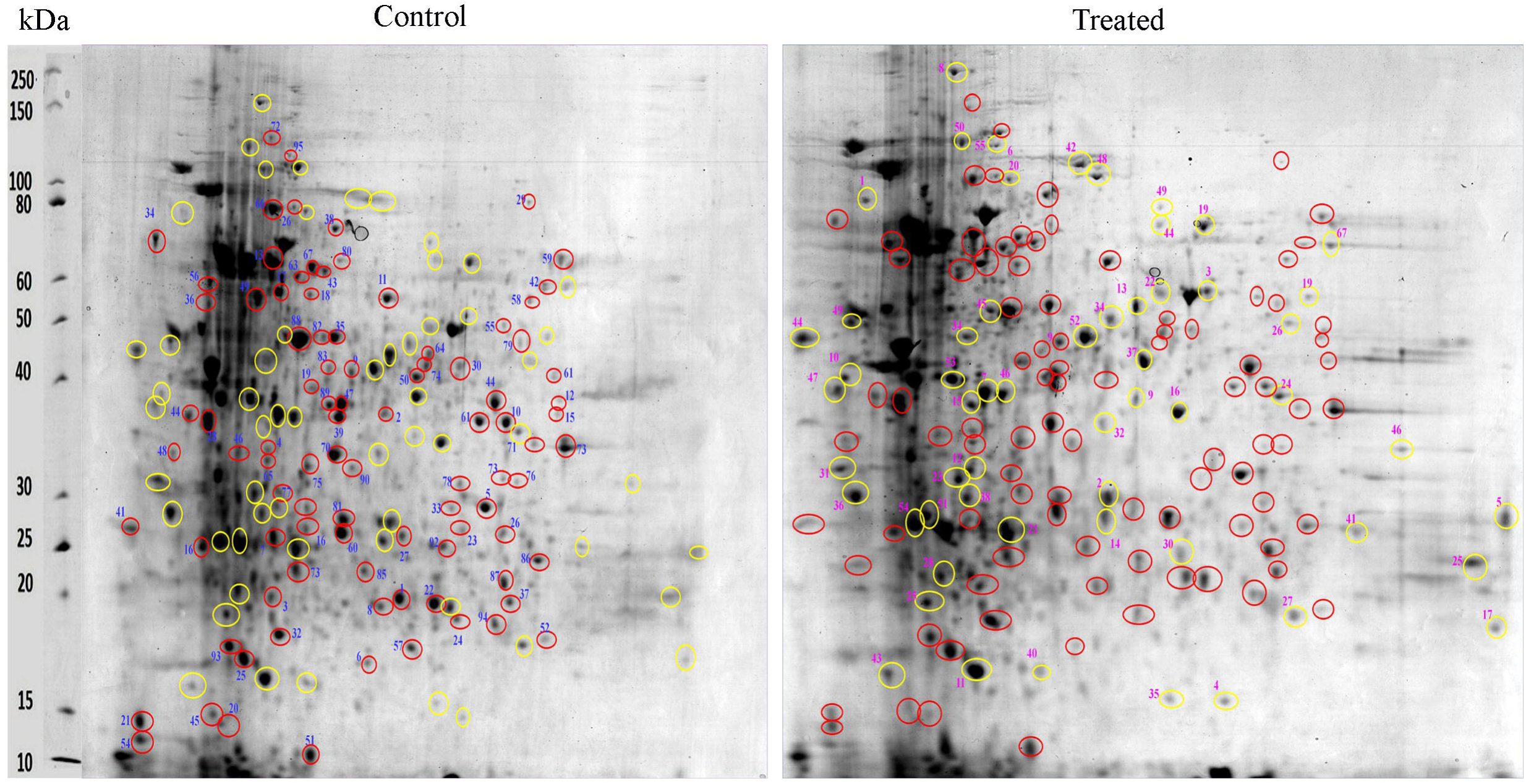
Gene Ontology analysis using the UniProtKB online tool showed that *C. elegans* regulatory proteins are involved in binding activity, catalytic activity, cell parts, cellular processes, metabolic processes, larval development, reproduction, and locomotion.

**Figure 4.**
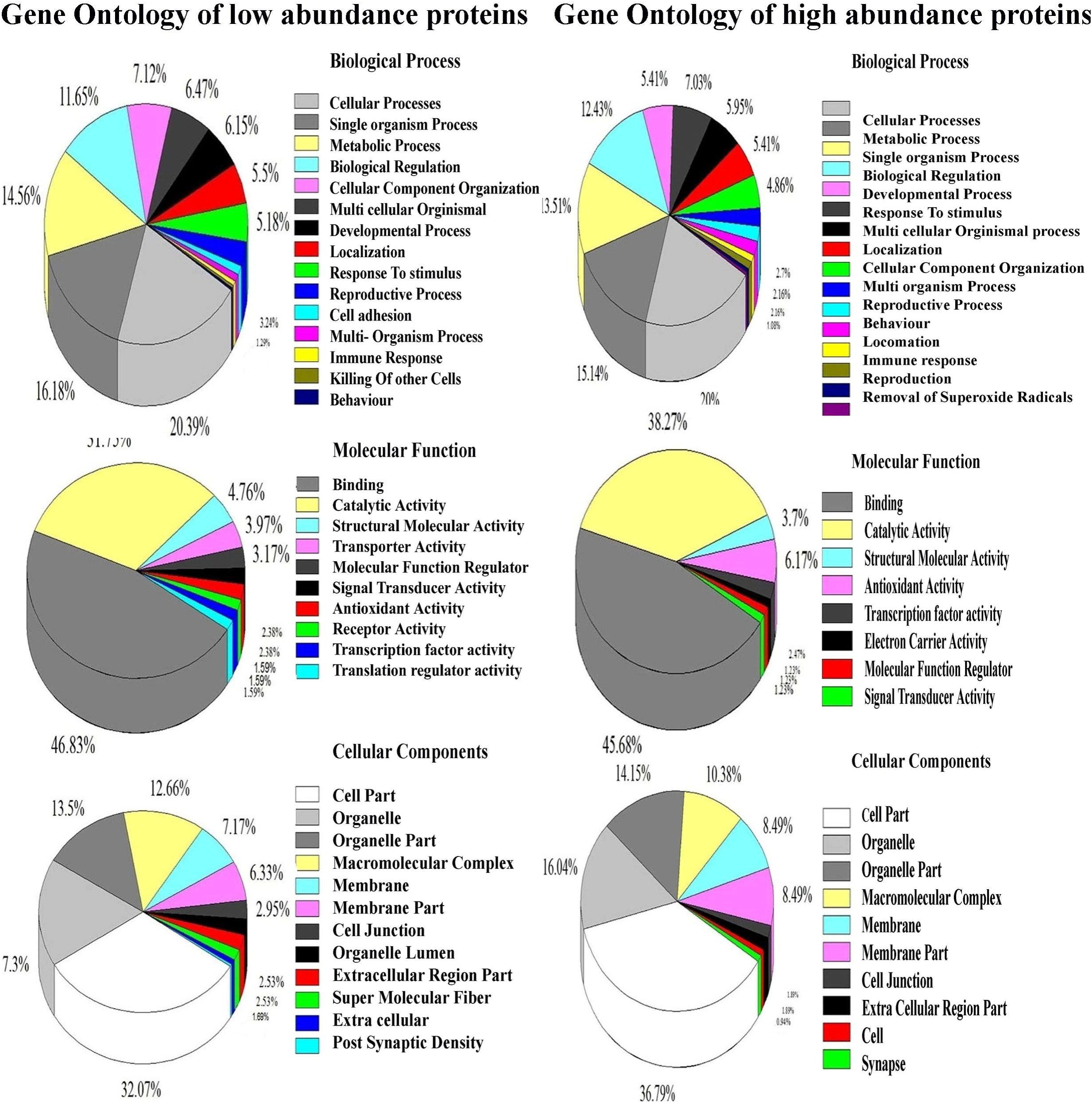
**A**) Interactome Map using the STRING tool with a medium confidence score (0.400), revealing an interaction between identified protein players which were regulated by 50% *S.* Typhi toxin protein exposure in *C. elegans*. **B**) Functional annotations and protein enrichment score of regulated proteins were performed using the DAVID tool. The biological functions which have the highest number of proteins regulated in response to *S.* Typhi protein fraction exposure are cytoskeleton, metal binding, cell adhesion protease and redox processes. **C)** Physiological function having highest protein enrichment score in responsible *S.* Typhi protein fraction exposure is embryonic development. **D**) Light microscopic images of *C. elegans* embryos treated to *S.* Typhi toxin proteome fraction (concentration 1.5 □ 1.0 mg/mL) showed reproductive failure and degenerated embryo formation.

### 2.3 *C. elegans* protein classes regulated by *S.* Typhi proteome

#### 2.3.1 Regulation of reproduction and embryo development proteins

The GO analysis showed 56 regulated proteins, which appears to play important role in embryogenesis and reproduction among which the proteins identified responsible for the embryonic development include LIN-28, UNC-60, SPL-1, ZYX, MUT, HMP, Y43F4A.1, STI-1, NPP-1, SAS-5, UNC-98, ZYG-1, PIG-1 CED-2 and TDO-2. Light microscopic images [Nikon Eclipse TI-s, Japan], of treated *C. elegans* showed reproductive failure and produced degenerated embryos compared with that of worms fed with *E. coli* OP50 (control) (**Figure 4D**). *C elegans* embryo cells use to terminally differentiate within 12 hrs of incubation at 20°C to enter life cycle events (L1 larvae stage). The effect of bacterial toxin protein on nematode embryogenesis was further confirmed by treating the isolated embryo cells with precipitated protein toxins (fractions 50% concentration, 1.5 □ 1.0 mg/mL) where it was found to fail enter into the L1 larvae stage. *C. elegans* embryos treated with toxins showed retarded growth and development; even after 48 hrs of incubation, none of the embryo enters into L1 larvae stage of animal (**Table 1**). However, isolated embryo cells from the control, entered into the normal L1 larvae developmental stage after 12 hrs of incubation. These experimental data showed that *S.* Typhi toxin protein exposure affects the *C. elegans* fertility, by directly retarding the embryogenesis. It is evident that increasing concentrations of toxin protein significantly (*p* < 0.05) decreased larvae stage formation.

#### 2.3.2 Regulation of oxidant and antioxidant proteins of *C. elegans*

Several proteins involved in oxidative stress were modulated during toxin exposure, indicating strong involvement in the cell response to toxins exposure. The measurement of extracellular ROS by DFC staining revealed that Typhi toxin protein treated N2 worms have elevated levels of ROS generation. ROS induction was examined at three-time points (12, 24, and 48 hrs) and it was found that ROS generation was high in treated worms compared with that of control samples (**Figure 5A**). Furthermore, the H_2_O_2_ production level was significantly (*p* < 0.05) higher in Typhi toxin protein treated samples compared with that of control for all tested time points provided in (**Figure 5B**). The high level of H_2_O_2_ indicated the role of reactive oxygen species for nematode mortality.

**Figure 5.**
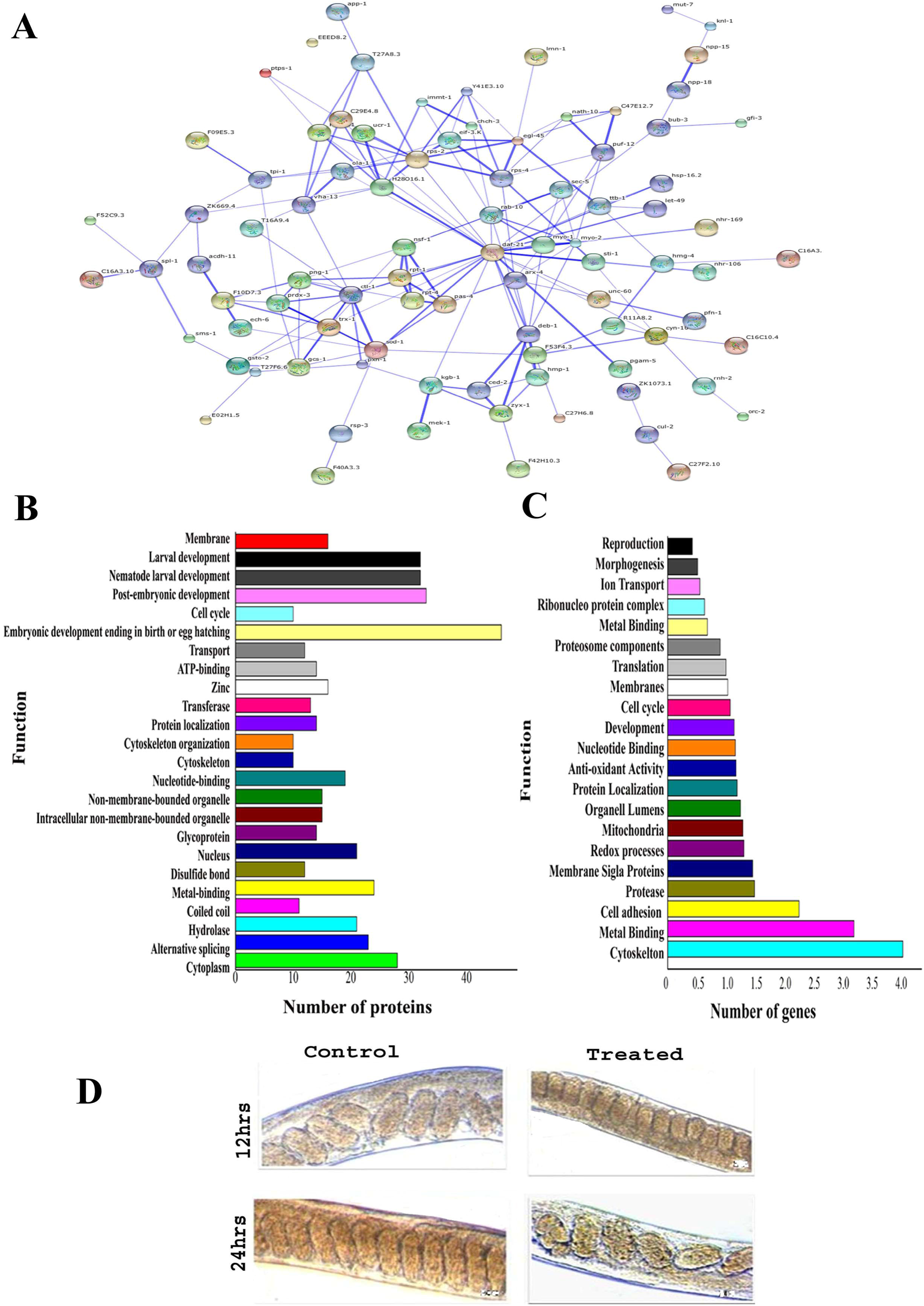
Quantitative analysis of oxidant and antioxidant proteins of *C. elegans* fed with *E. coli* OP50 and treated with *S.* Typhi toxin proteins and *E. coli* OP50 total proteins. **A**) ROS estimation, **B**) H_2_O_2_ estimation, **C**) Quantification of SOD activity, **D**) Quantification of catalase activity and **E**) Protein carbonyl content estimation. Data were expressed as mean value of three experiments and the error bars represent SD ± mean (\**p* < 0.05).

Several upregulated antioxidant proteins viz, superoxide dismutase enzyme (SOD), catalase (CTL), peroxiredoxin (PRDX), peroxidise (SKPO-1), thioredoxin (TRX) and glutamate-cystinease (GCS) have corroborated the *in vivo* detection of H_2_O_2_ associated ROS generation that directly leads to accumulation of molecular damage. The estimation of SOD was evaluated in both control and treated samples at three different time points (12, 24, and 48 hrs). The measurement of SOD alone showed significant increase of 10 fold in Typhi toxin protein treated L4 stage animals compared with that of control provided in (**Figure 5C**). Quantitative spectrophotometric analysis of catalase activity of L4 stage treated nematodes showed significantly (*p* < 0.05) high catalase activity for 12 and 24 hrs time points compared with that of control provided in (**Figure 5D**). In contrast, treated *C. elegans* at 48 hrs showed significantly (*p* < 0.05) decreased catalase activity, which suggested the rescue against H_2_O_2_ free radicals and other oxidative stresses appear to be decreased. In host cells, protein carbonyls content were also measured to determine the oxidative damage in the worms. The estimation of protein carbonyl contents were evaluated in both control and treated *C. elegans* at 12, 24, and 48 hrs. At three time points compared with that of control (1.83333 ± 1.2 nM/mg, 2.3948 ± 1.15 nM/mg and 4.87308 ± 1.10nM/mg), treated worms showed significantly (*p* < 0.05) increased carbonyl content level (2.936253 ± 1.20 nM/mg, 11.78944 ± 1.20 nM/mg and 16.04382 ± 1.25 nM/mg) at 12, 24 and 48 hrs, respectively (**Figure 5E**). Elevated levels of protein carbonyls could be caused by an increase in protein oxidation or by a decrease in the turnover rate of the oxidized proteins in treated sample.

#### 2.3.3 Regulation of chaperones and stress proteins

Several chaperones (DAF-21(HSP-90), HSP-16.2, GCS-1, STI-1, GSTO-2, PXN-1, TRX-1) have been differentially regulated in treated *C. elegans*. STRING analysis showed direct interaction of HSP proteins with antioxidant (SOD-1, SKPO-1, CTL-1, PRDX-3, SKPO-1, GCS-1) and stress (STI-1 and TRX-1) proteins. Defence against oxidative stress is very much interrelated with network of heat shock or stress-proteins (Espinosa-Diez *et al*., 2015). To validate the MALDI-TOF/TOF/MS results, the expression of DAF-21 protein was evaluated using Western blotting. The Western blotting results showed downregulation of DAF-21 protein in treated samples compared with that of controls provided in (**Figure 6A**). The 3D view of DAF-21 protein (**Figure 6B**). STRING analysis of DAF-21 protein showed direct interaction with several MAP Kinase pathway specific proteins (**Figure 6C**). Western blotting analysis of these candidates MAP Kinase pathway specific proteins (JNK-1, p38, SGK-1 and HSF-1) were also downregulated compared with that of controls provided in (**Figure 6A**).

**Figure 6.**
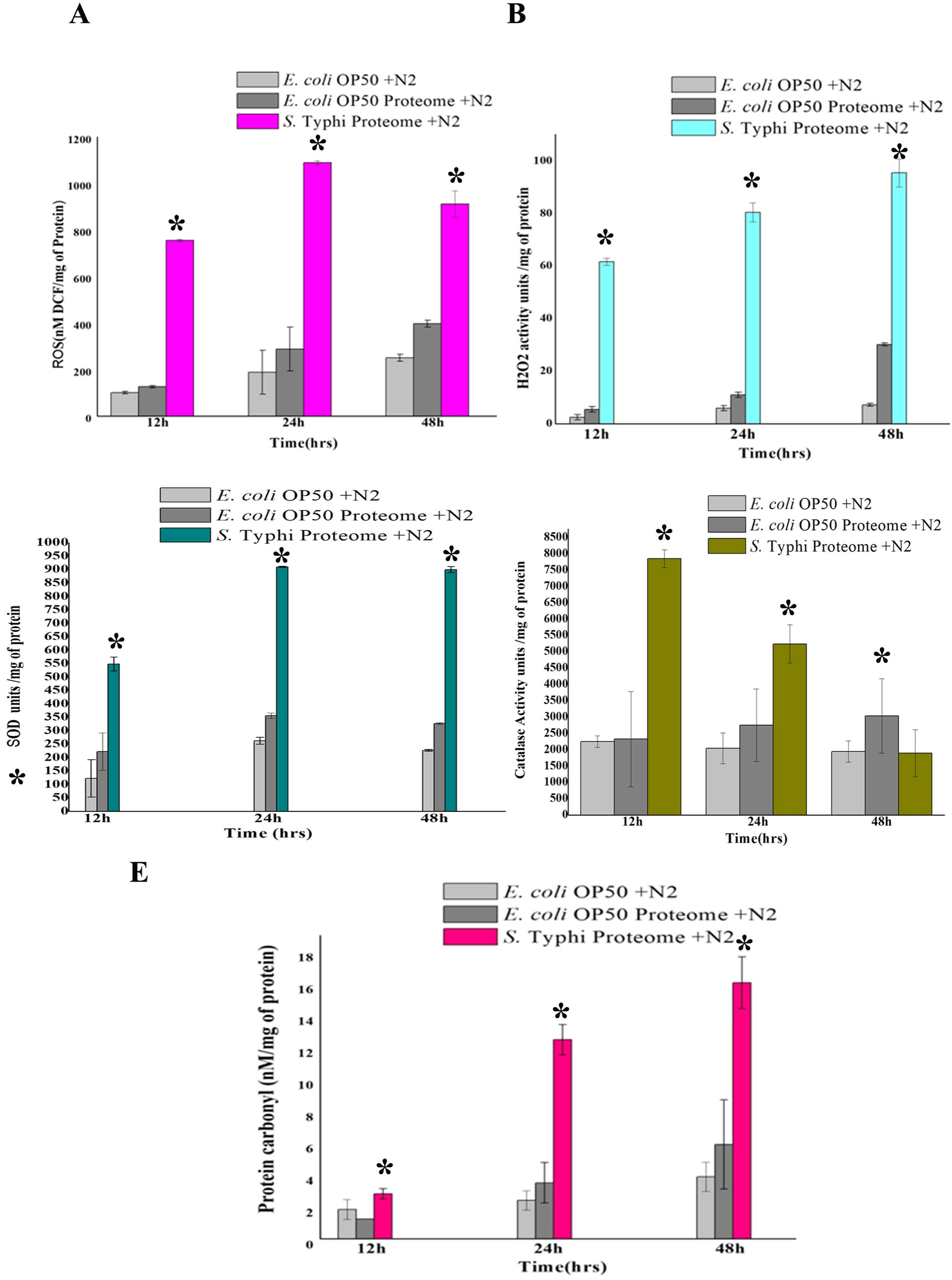
**A**) Western blotting analysis showed downregulation of JNK-1, p38, HSP-90, SGK-1 and HSP-1 at 12 and 24 hrs, detected by using the specific antibodies. The protein expression levels at each time point were normalized with β-actin. **B**) Proteins analysed by Western blotting showed a close functional regulation interactions with each other identified using the STRING tool. **C**) DIA analysis represented 3D view of regulated DAF-21 protein *Differences were considered significant at *p* < 0.05.

#### 2.3.4 Regulation of lipid metabolism proteins

Proteins related to lipid metabolism identified, in this study include ADS-1, ZK669.4, SMS-1, BRE-4 and PNG-1. A regulation of these proteins indicated an altered fat storage in living organism. To validate the fat and lipid deposition change in treated *C. elegans*, Oil-Red-O staining was performed. *C. elegans* stained with yellow gold fluorescence are lipid droplets and red fluorescence is phospholipids. Microscopic images reveal the difference in fat storage. A decrease in red staining was observed in treated *C. elegans* compared with that of control. This result suggested the impact of toxin protein on the level of fat molecules in a host system (**Figure 7A**). *C. elegans* treated with *S.* Typhi proteome exhibited significant ROS generation provided in (**Figure 7A)**. Quantifying a lipid peroxidation is an effective way to measure the effect of oxidative damage. Concurrently estimation of *C. elegans* lipid peroxidation for both control and treated at three time points (12, 24, and 48 hrs) showed significantly increased level of TBARS (2-thiobarbituric acid-reactive substances) formation by H_2_O_2,_ which induced lipid peroxidation in treated samples compared with of control (**Figure 7B**).

**Figure 7.**
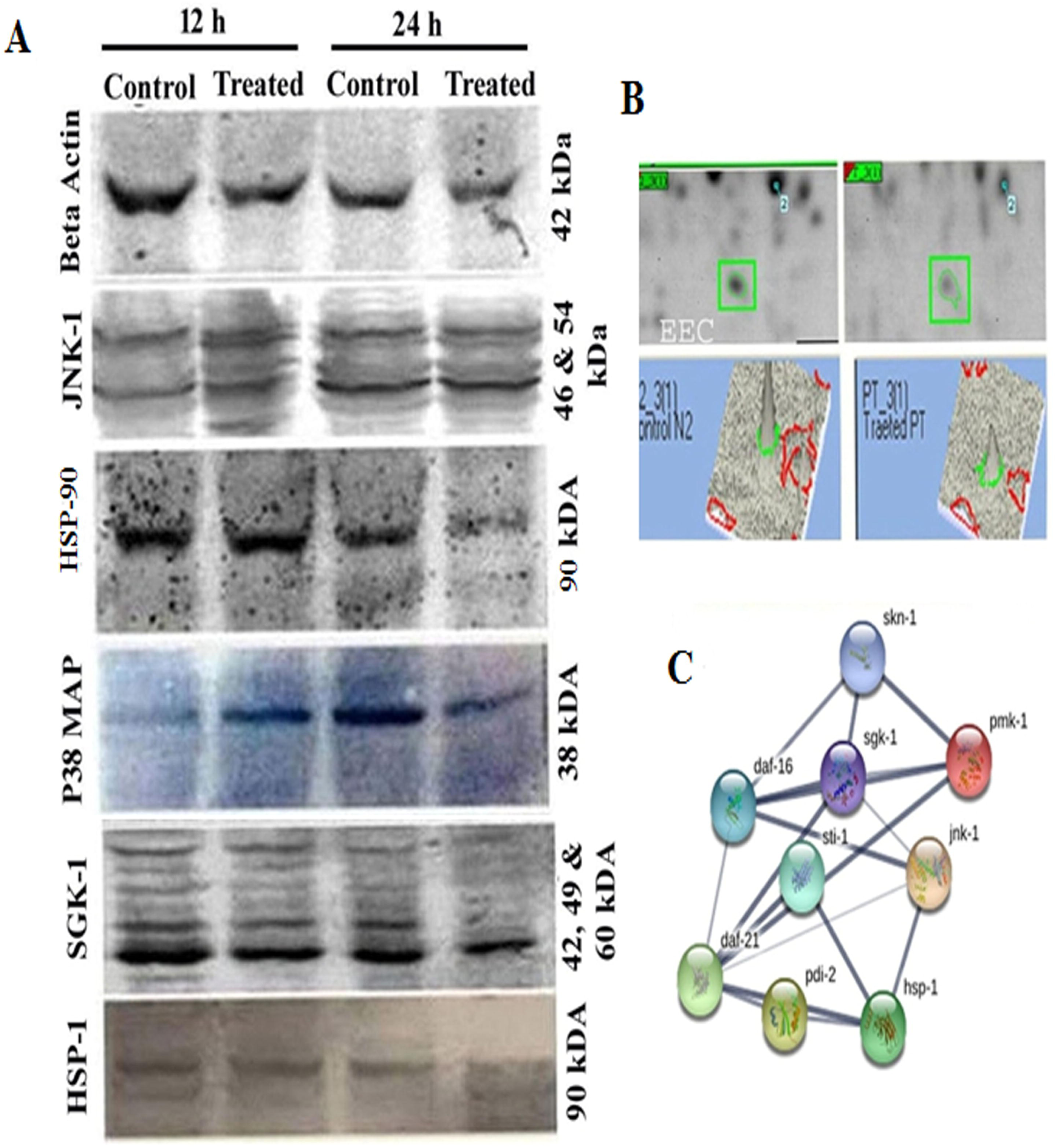
**A**) Control N2 and *S.* Typhi proteome treated worms were stained with Oil-Red-O reagent. **B**) Lipid peroxidation quantification. *Differences were considered significant at *p* < 0.05.

#### 2.3.5 Regulation of ATP energy production proteins

The proteins related to energy metabolism, identified in this study include VHA-13, H28O16.1, OLA-1, MAI-1, LET-754, C29E4.8 and F40F8.1. The VHA-13 encodes V-ATPase; H28O16.1 encodes alpha subunit of mitochondrial ATP synthase. All these enzymes are responsible for aerobic respiration, which is the efficient pathway for metabolic energy. The downregulation of these enzymes likely exhibited an energy deficit which appeared to be one of the reasons for *C. elegans* death. Hence, the total ATP production of *C. elegans* was measured, where the intracellular ATP level was significantly (*p* < 0.05) low in treated samples compared to control, (**Figure 8**). This result indicated that the expression of the ATP production in worms have decreased or inhibited due to intoxication which was corroborated with the expression levels of ATP generating proteins in this study.

**Figure 8.**
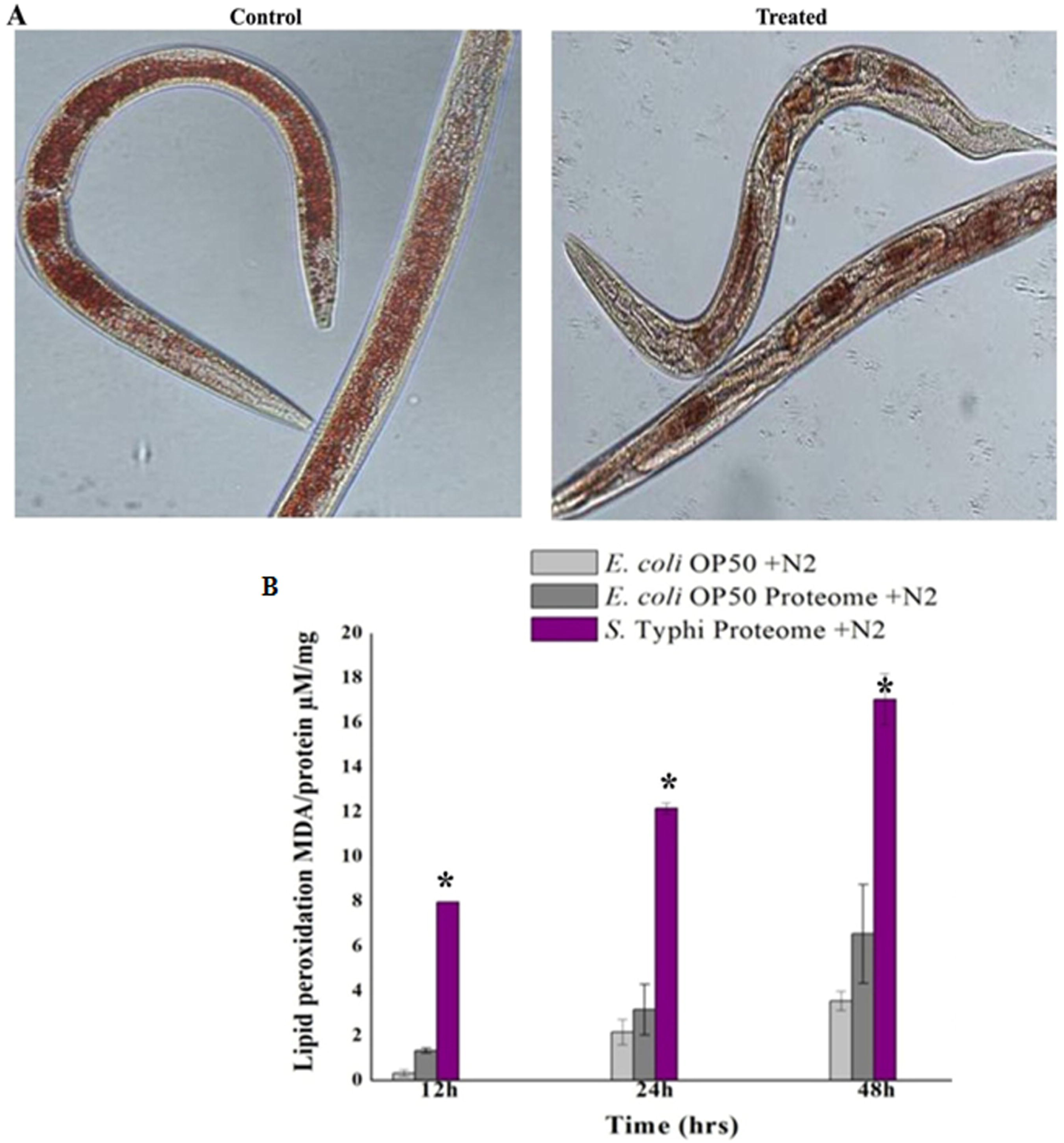
Effects of *S.* Typhi toxin proteins on the energy metabolism. Estimation of the total intracellular ATP in *C. elegans* was significantly (\**p* < 0.05) reduced in treated samples compared with that of controls (fed with *E. coli* OP50).

#### 2.3.6 Regulation of cytoskeletal organization proteins

After 12 hrs of treatment with toxin protein, worms were transferred to NGM plates (food source) as represented in (**Figure 9A**). The worm treated to 500 μg/mL of *S.* Typhi toxin proteome was active and precede significantly (*p* < 0.05) a normal life span, there body movements and locomotion was also normal. However, worms treated to 1 mg/mL of toxin proteins exhibited a significantly (*p* < 0.05) a mean life span of 60 ± 5 hrs with retarded locomotion and body movements. The worms treated to 1.5 mg/mL of toxin proteins were immobile, produced no progeny and exhibited significantly (*p* < 0.05) a mean life span of 30 ± 5 hrs as provided in (**Figure 9B**).

**Figure 9.**
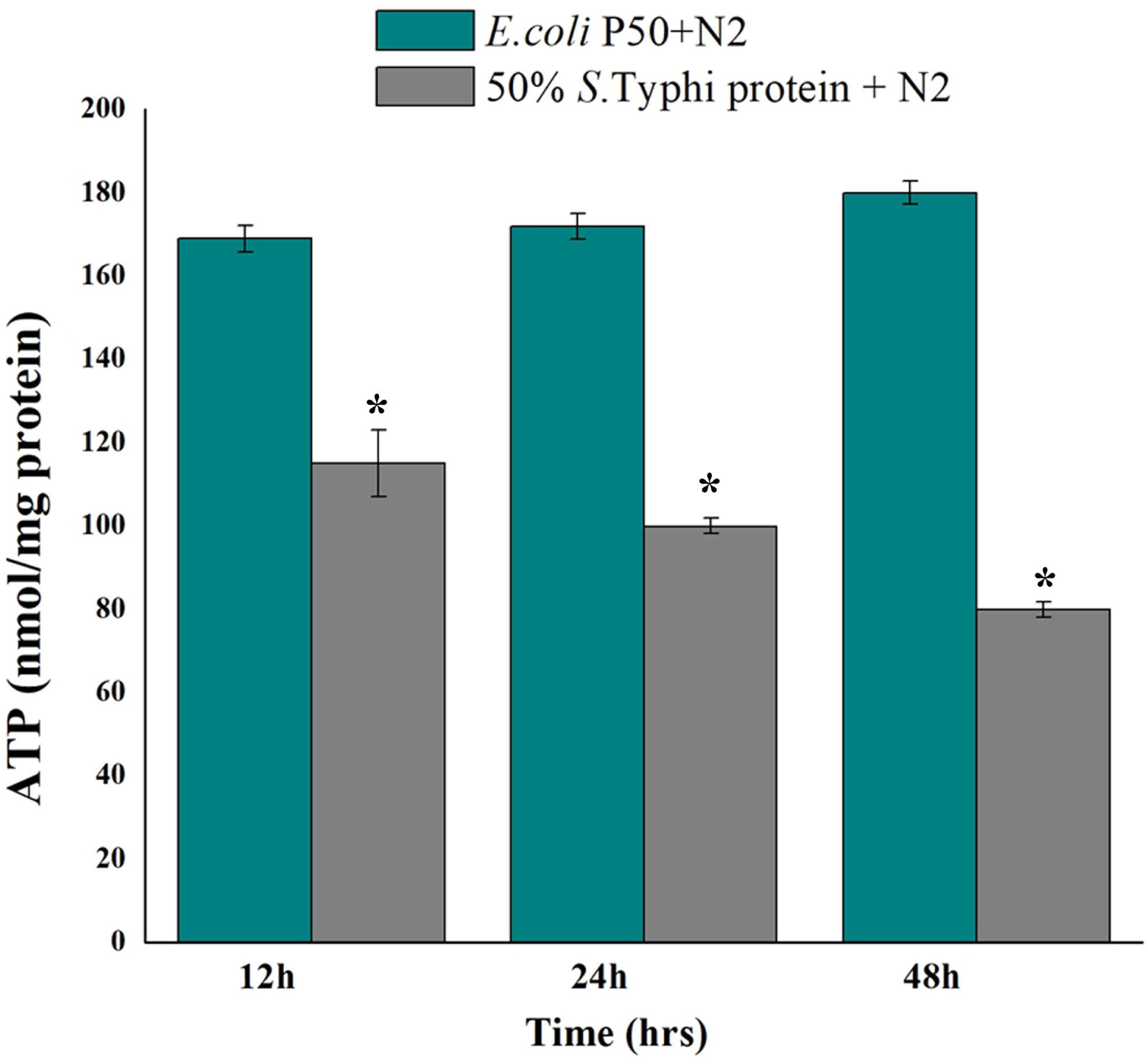
**A**) Graphical representation of behavioural assa**y B**) Impact of *E. coli* OP50 food source to worms pre-treated 12 hrs with *S.* Typhi toxin protein fraction. Worms treated with various concentrations of precipitated proteins showed varied physiological characteristic and life span.. **C**) Microscopic imaging of *S.* Typhi toxin proteins treated *myo-*2 and *col-*19 GFP-tagged strain showed high level of fluorescence compared to controls. *Differences were considered significant at *p* < 0.05.

Also, in this study, several identified proteins viz., UNC-54/60/98, MYO -1/2, LMN-1, ZYX-1, ARX-4, IFTA-2 and DEB-1 which are crucial for locomotion and muscle contraction was downregulated. Interestingly, STRING analysis showed close association between these genes as provided in (**Figure 4A**). UNC-54 is the myosin heavy chain protein which is essentially expressed in *C. elegans* for locomotion and egg-laying. The downregulated protein MYO-2 and COL-19 were confirmed by the specific *C. elegans* based experiments. The microscopic imaging studies revealed that treated *myo-*2 and *col-*19-GFP tagged protein strains showed high fluorescence compared to control strains fed with *E. coli* OP50 (**Figure 9C**). An overview of proteins and pathways targeted by *S.* Typhi toxin proteome in *C. elegans* during exposure are presented in **Figure 10.4.**

**Figure 10.**
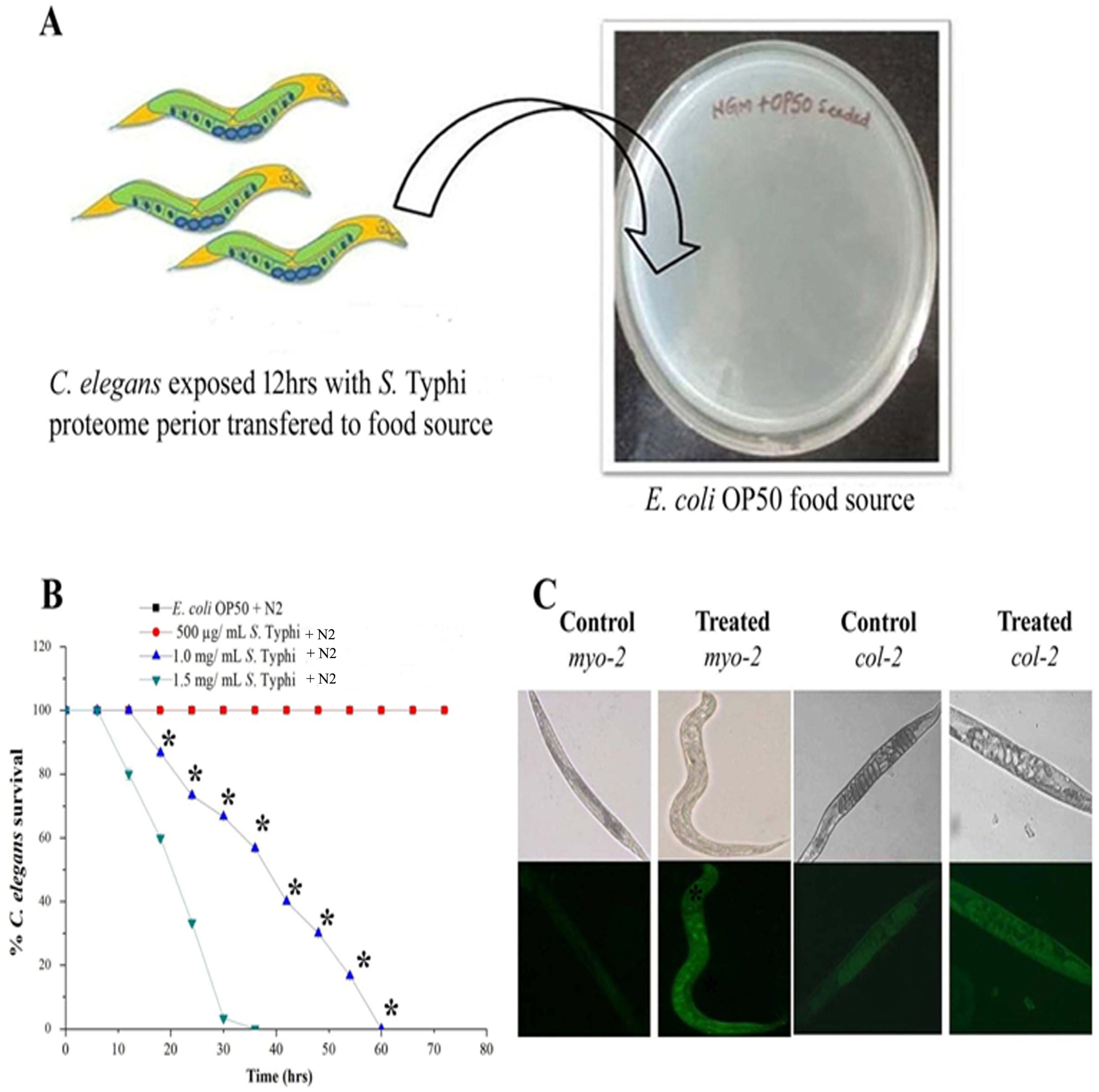
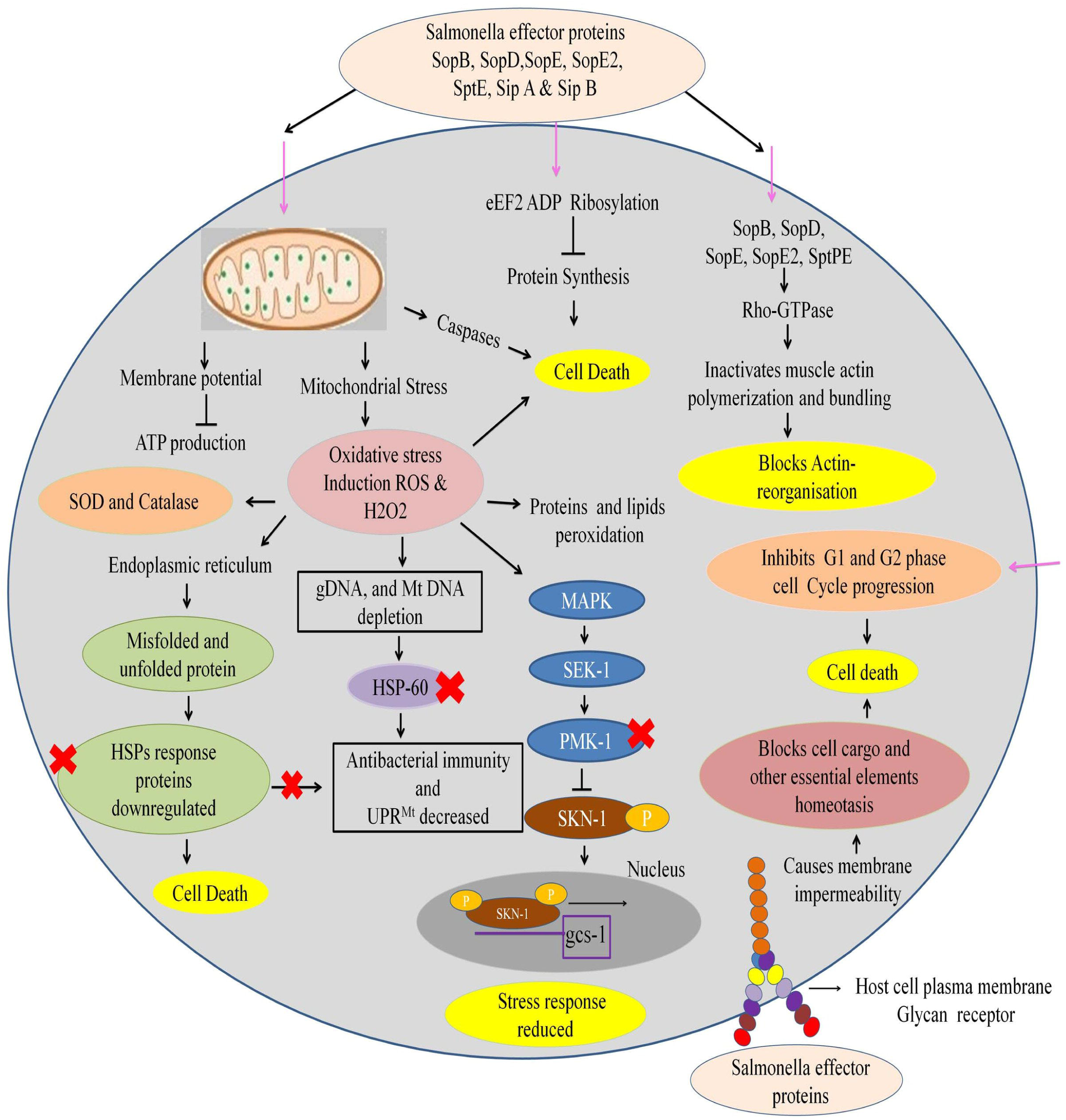
An overview of proteins and pathways activated and targeted by *S.* Typhi toxin in *C. elegans* are presented in this figure. The Salmonella effector proteins SopB, SopD, SopE, SopE2, SipA & Sip B enter into the host cell cytosol. These complex macromolecules employ several mechanisms in a highly regulated manner and manipulate the host cell in various ways. The toxic macromolecules inhibit protein synthesis by ADP-Ribosylation process. Some of these toxic macromolecules act on membrane voltage gated channels (VADC) causing depolarisation of mitochondrial potential. The VADC blockage inhibits the ADP, Pi and pyruvate to cross into the mitochondrial membranes to generate ATP. The mitochondrial stress generates Reactive oxygen species (ROS). Salmonella effector proteins reduces the antioxidant defence mechanism of host and makes it immune compromised. HSPs promote the folding of the imported protein to its native conformation. Downregulation of HSPs protein in treated samples might have increased the unfolded proteins in host. HSP-60 is important molecular players in *C. elegans* involved in the activation of the MAP Kinase and UPR^Mt^. Functional loss of HSP-60 regulatory protein causes the downregulation of the MAP Kinase pathway which might be the reason of nematode susceptibility

## 3. Discussion

The nematode model, *C. elegans* is well suited for the studies including developmental biology, molecular biology, host-pathogen interactions, neurobiology and hypoxia (Schouest *et al*., 2009; Marsh *et al*., 2012; Ren *et al*., 2010; Kamaladevi *et al*., 2017). The present study highlighted the impact of bacterial whole proteome on *C. elegans* with the specific attention to proteomic alternations induced by the same. This study showed that *S.* Typhi proteins are significantly involved in differential regulation of *C. elegans* host proteome. Based on the densitometry analysis of 2D-GE images, among 477 spots found 95 and 55 spots were significantly downregulated and upregulated, respectively. The combined spectra of MALDI-TOF/TOF/MS were searched against the Swiss-Prot database of *C. elegans* using a MASCOT engine for identification and characterization. The regulated proteins were identified and annotated for their specific biological and molecular functions.

*C. elegans* regulates a diversified molecular response upon adverse environmental conditions, bacterial infections and physiological stress to promote adaptation for survival. In this study, intoxication has regulated the expression level of HSP (HSP-90, HSP-16.2, HSP-6 and HSP-4) in wild-type N2 worms. This is corroborated with the previous reports (Frydman *et al*., 2001; Prithika *et al*., 2016) where it is stated that HSPs provide an immediate response during stress, tissue damage or bacterial infection It is anticipated that the identified regulation of HSP during S. Typhi may act by modulating structurally denatured/ misfolded proteins to retain their native confirmation and degrading the proteins which are not properly refolded as described earlier (Soti *et al*., 2005; Powers *et al*., 2010). Many reports are also in agreement about the role of HSP in bacterial infection and immunity (Wang *et al*., 2017; Prithika *et al*., 2016, JebaMercy *et al*., 2016; Durai *et al*., 2014). Collectively, our results suggested that HSP played a vital role in the stress response of *C. elegans* against *S.* Typhi proteome. Furthermore, the study have found the downregulation of protein DAF-21 (HSP-90) and its interaction with proteins including muscle contraction (MYO-1/2), locomotion (UNC-54), energy hydrolysis (LET-754), ATP synthase complex (H28O16.1), stress response (STI-1), serine/threonine kinase activity (KGB-1), oxidoreductase activity (TRX-1) and HSP-16.2 (**Figure 4A**). There are certain earlier studies in support of the above finding, which have suggested the important role of DAF-21 in increasing the *C. elegans* immunity against bacterial pathogenesis (Mohri-Shiomi *et al*., 2008). In addition, the DAF-21 played a key role in the regulation of MAP Kinase pathway by phosphorylating the mitogen activated protein kinase (MPK-1) (Green *et al*., 2011). In this study, DAF-21 protein showed a significant fold change compared with that of control (**Figure 6B**). Since, DAF-21 plays a crucial role in regulating MAP Kinase pathway (Green *et al*., 2011), the study have explored its associated proteins (JNK-1, SGK-1, PDI-1, p38 and HSP-1) by western blot analysis (**Figure 6A**). In addition it is probable that functional loss of DAF-21 might have inhibited MAP Kinase pathway which in turn might be the cause for nematode susceptibility DAF-2 was appeared to be important molecular player that activates MAP Kinase for rescuing host from toxins (Green *et al*., 2011). The expression level of antioxidant enzyme verified the oxidative damage in exposed worms. These findings have corroborated the *in vivo* detection of H_2_O_2_ and ROS generation which directly leads to accumulation of molecular damage in the host cells. Furthermore, upregulated glutathione transferase enzyme families may be involved in xenobiotic detoxification which provided resistance against pathogen in *C. elegans* as described earlier (Lindblom *et al*., 2006).

*C. elegans* fat molecule is dynamic in nature; it increases both in size and number during development (Hellerer *et al*., 2007). In our study, regulation of several molecular players related to lipid metabolism [ADS-1, ZK669.4, SMS-1, BRE-4 and PNG-1] in response to toxin exposure was identified. This suggested that alteration in lipid metabolism occurs in response to intoxication. The ADS-1 (Alkyl-Dihydroxyacetone phosphate synthase) protein is an ortholog of human AGPS (Alkylglyceron phosphate synthase). ADS-1 protein is required for the initial stage of ether lipid biosynthetic pathway (Shi *et al*., 2016) which is required for the initial stage of ether lipid biosynthetic pathway (Shi *et al*., 2016). The downregulation of ADS-1 was found in our study which indicates the inhibition of lipid biosynthesis during exposure. Humans born with mutations in *agps* gene die early because of severe growth and neurological defects (Braverman *et al*., 2012). The bre-4 gene encodes β-1, 4-N-acetylgalactosaminyl transferase is required for the toxicity of Bacillus thuringiensis Cry5B toxin protein (Kho *et al*., 2011). The galactose-β 1,4-N-acetylglucosamine containing carbohydrate chains are attached with proteins and lipids that binds with the galectins group (Kasai *et al*., 1996), plays an important role in biological events such as development, immunity and cancer defense (Yang *et al*., 2008; Boscher *et al*., 2011). The downregulation of *C. elegans* fat and lipid metabolism proteins appeared to be directly responsible for reduced levels of stored lipids and fatty acids. Oil-Red-O staining of nematodes produced visual patterns of neutral lipids that are representative of biochemical determinations of fat levels (Wahlby *et al*., 2014; Fouad *et al*., 2014). The fatty acids play a critical role in modulating lipid/yolk level in the oocytes and regulating reproductive efficiency of *C. elegans* (Chen *et al*., 2016). In current investigation the significant changes in lipid metabolisms have proven to be closely related with hypersensitivity of the worms towards *S.* Typhi toxin proteins.

Few candidate molecular players namely, LIN-28, UNC-60, SPL-1, MUT, HMP, Y43F4A.1, STI-1, NPP-1, SAS-5, UNC-98, ZYG-1, PIG-1 CED-2 and TDO-2 were regulated during exposure to toxins which are necessary for reproductive events and oocytes development [www.wormbase.com]. Toxin proteins subsequently affected the reproductive events of *C. elegans* whereas control worms didn’t showed any egg laying defects. In fact, the downregulation of molecular players might be the reason to induce the morphologically degenerated and developmentally abnormal embryos. The downregulation of ATP during toxin protein exposure suggested that the host probably had an energy deficit which could have lead to *C. elegans* mortality.

During the bacterial proteome exposure, it was observed that UNC-54/60/98, MYO-1/2, LMN-1, ZYX-1, ARX-4, and DEB-1 proteins were downregulated which are necessary for muscle myosin filament assembly, locomotion, depolymerisation, contraction and regulation of actin polymerization. DIM-1 is an immunoglobulin protein essential for maintaining body wall muscle integrity (Rogalski *et al*., 2003) and localizes in between the dense bodies and the region of the muscle cell membrane (Otey *et al*., 2005). The DIM-1 can alleviate the locomotion defects caused by toxins (Wang *et al*., 2017). Regulation of DIM-1 protein directly affects the expression of UNC-54, ZYX-1 and DEB-1 proteins and reduction of these proteins causes severe muscle disruption and paralysis (Etheridge *et al*., 2012; Rogalski *et al*., 2003). The protein ZYX-1(Zyxin) acts as muscle mechanical stabilizer and a sensor for muscle cell damage (Lecroisey *et al*., 2013). The ZYX-1 protein regulates, stabilizes and maintains posterior mechanosensory neuron extension, new synapse formation and growth during larval development (Luo *et al*., 2014). Our finding supported the role and involvement of the above players during toxin proteome exposure.. Several identified regulated proteins are involved in Na^2+^ and Ca^2+^ voltage gated channels which lead to degradation in the synapses of neurons, immune response pathway and normal cellular process. In addition few uncharacterized proteins (T19C3.6, TPI-1, F52C9.3, F26E4.3, EEED8.2, E02H1.5, F10.D7.3, R166.3 and C05D10.4) were also found to be regulated.

The inadequacy and the high costs associated with mammalian testing reduced the possibility to evaluate the toxicity of a variety of bacterial toxins, environmental chemicals and pollutants. Our study attests the utility of *C. elegans* as an emerging model in toxicological sciences. Our findings reveal that the amount of oxidized proteins increases in several folds which require a highly organised participation of chaperones to rectify the damage in protein conformational damage. These changes can also occur during aging. In contrast, enhanced antioxidant systems as well as over expression of heat shock proteins lead to longevity. Taken together, our data suggest that altered HSP-90 protein expression, MAPK, JNK signal pathways, ZNK-1, BRE-4 and BRE-5 were observed for the first time to participate in *C. elegans* defence mechanism against *S.* Typhi proteome, although their detailed functions and mechanisms in stress responses remain ambiguous. This information will help broaden our knowledge on the mechanism of host-toxin interaction.

## 4. Materials and Methods

### 4.1 Maintenance of nematode, *C. elegans* and bacterial strains

Nematodes used in this study were wild-type N2 Bristol and out crossed mutants, *myo*-2 (EG5568), *col-*19 (TP12), obtained from the CGC (Caenorhabditis Genetic Centre). Synchronized L4 stage worms used for *in vivo* exposure experiments were obtained by bleaching, as described previously Theresa Stiernagle (2006). The L4 stage worms were washed with M9 buffer (3g KH_2_PO_4_, 6g Na_2_HPO_4_, 5g NaCl and 1 mL of 1 M MgSO_4_ to 1000 mL H_2_O sterilized by autoclaving) and used for all the *in vivo* exposure experiments. All physiological assays including short time exposure were performed using isolated toxin proteins of *S. enteric serovar* Typhi (MTCC 733), which was purchased from Microbial Type Culture Collection and Gene Bank (MTCC). All strains were maintained frozen at -80°C in the peptone solution (2% peptone, 5% glycerol). The frozen bacteria were usually grown on LB plates with 1.5% agar at 37°C. A single colony picked from the plate was inoculated into 3 mL LB medium, and then the overnight culture was diluted 1:20 into 300 mL LB broth (with 0.3 M NaCl to increase bacterial invasion).

### 4.2 Ammonium sulphate (AS) fractionation

The soluble proteins from the *S.* Typhi and *E. coli* OP50 bacteria pellets were obtained by suspending the microbial cell pellets in 1X PBS (8g NaCl, 0.2g KCl, 1.44g Na_2_HPO_4_ and 0.44g KH_2_PO_4_, pH 7.5), disrupted with an ultrasonicator (Sonics & Materials, Danbury, CT, USA) and centrifuged at 17000 × g for 20 min at 4°C. Before precipitation, the supernatant was filtered by using 0.2 micron filter paper. *S.* Typhi whole-cell extracted proteins were enriched by (NH_4_)_2_SO_4_ precipitation for concentrating the protein samples as per (Wingfield *et al.*, 2001., Park *et al*., 2008) and also the EnCor Biotechnology Inc (https://www.encorbio.com/protocols/AM-SO4) method which conveniently determined the amount of solid ammonium sulphate required to reach a given saturation. The precipitated protein fractions were desalted by high performance centrifuge Zeba spin desalted column as per manufacturers protocol. The toxicity assays including quantitative growth, embryonic development, oxidant and antioxidant assays and lethal dose assays were performed in liquid media. The Ammonium nitrogen in solution was estimated by phenate method as per Park *et al;* 2009 with the assay medium contains 0.025 μg/mL of Ammonium nitrogen.

### 4.3 Physiological studies during exposure with *S.* Typhi toxin proteins in *C. elegans*

#### 4.3.1 Nematode liquid killing assay

Nematode killing assay were performed to determine the pathogenicity of *S.* Typhi proteome. Approximately, 20 L4 stage age-synchronized *C. elegans* were transferred from a lawn of *E. coli* OP50 to a 24-well plate containing *E. coli* OP50 (control) or isolated total proteins of *E. coli* OP50 (negative control) or isolated toxin proteins of *S.* Typhi (treated). The plates were incubated at 20°C and scored for viability of *C. elegans* every 6 hrs. Worms were considered dead upon failure to respond upon gentle touch using a worm picker containing platinum wire on the solid NGM plates. All the experiments were carried out in triplicates. Kaplan-Meier survival analysis was used to compare the mean lifespan of control and treated nematodes. The experiment was performed in biological triplicate, and the error bars represent the mean ± SD (\**p* < 0.05).

#### 4.3.2 *C. elegans* protein sample preparations

*C. elegans* treated to *S.* Typhi toxin proteins and *E. coli* OP50 bacteria for 24 hrs. After exposure, the worms were washed thoroughly with M9 buffer to take away the surface bound proteins and bacteria. The washed worms were immersed in 50 mM Tris-HCl buffer (pH 8.5 along with protease inhibitor cocktail) then sonicated on ice for 3 min at 10 second pulse interval, debris were removed by centrifugation at 7000 × g for 5 min and subsequently, the resulting supernatant was collected and purified using a 2D cleanup kit (GE Healthcare) as per manufacturer’s protocol. The Protein concentration was determined using Bradford reagent (Sigma Aldrich) (Bradford *et al*., 1976) and the protein concentration was maintained at 1mg per sample. The samples were dissolved in (7 M urea, 2 M thiourea, 4% CHAPS, 40 mM Dithiothreitol (DTT) and carrier ampholytes 2% v/v added (per manufactures protocol) and subjected to 2D-GE.

#### 4.3.3 Iso-electric focusing (IEF) and 2D-GE

During IEF (first dimension), proteins were separated based on their isoelectric point on 18-cm immobilized pH gradient (IPG) gel strips of pH 3-10 (GE Healthcare). Subsequent to overnight rehydration, IPG strips were subjected to IEF at 20°C under mineral oil with the following conditions: 3 hrs at 100V; 1 h at 500V; 1 hrs at 1000V; 2 hrs at 1000-5000V (gradient); 1 h at 5000V; 3 hrs at 5000-10000V (gradient) and final focussing was done for 2 hrs at 10000V. The current was set to 75 μA per IPG strip. Prior to SDS-PAGE, IPG strips were immersed twice for 15 min in equilibration buffer I [6 M urea, 30% (w/v) glycerol, 2% (w/v) sodium dodecyl sulphate (SDS) and 1% (w/v) DTT in 50 mM Tris-HCl buffer, pH 8.8] followed by equilibration buffer II [6 M urea, 30% (w/v) glycerol, 2% (w/v) SDS and 2.5% (w/v) iodacetamide (IAA) in 50 mM Tris-HCl buffer, pH 8.8]. After equilibration, proteins were separated based on their molecular weight using 12.5% SDS-PAGE (second dimension). Electrophoresis was performed at 100V (200 mA) for 1 h and 150V (300 mA) for 7-8 hrs in Ettan DALT six apparatus (GE Healthcare). After electrophoresis, gels were kept in fixative solution (containing 40% methanol, 10% glacial acetic acid and 50% Milli Q H_2_O) for overnight and washed thrice with Milli-Q H_2_O for 20 min each. Protein spots were stained by colloidal coomassie brilliant blue (CBB) G-250 staining solution (containing 10% ortho phosphoric acid, 10% ammonium sulphate, 20% methanol and 0.12% CBB) for 12 hrs on rotary agitator. Subsequently, the gels were destained with Milli-Q H_2_O for 4 hrs to reduce the background noise.

#### 4.3.4 Trypsin digestion of differentially regulated protein spots and mass spectrometric analysis

Destained gels were scanned with a densitometry image scanner (Image Scanner III, GE Healthcare) at 300 dpi resolution and captured using Lab Scan 6.0 software. The raw images were analysed using Image Master 2D Platinum 7 (GE Health care). Based on the densitometry analysis of gel images which detected about total 898 protein spots amongst, 477 spots were satisfied the arbitrary parameters in control and treated gels. Interested spots with more than 1.5 fold changes in the intensity were excised manually. Subsequently, prior to in-gel trypsin digestion, the excised proteins were completely destained by washing with destaining solution (containing 50% acetonitrile and 25 mM ammonium bicarbonate) and dehydrated in 100% acetonitrile (ACN) for 10 min, then dried under vacuum for 30 min. After reduction and alkylation as per the standard protocol, in-gel trypsinisation was performed with 5 μL of trypsin buffer (10 mM NH_4_HCO_3_ in 10% ACN) containing 80 ng of trypsin (Sigma Aldrich) and incubated at 37°C for 16 hrs. After incubation, peptides were extracted by 0.1% trifluoroacetic acid (TFA) in 60% ACN by bath sonication (10 min) subsequently dehydrated by 100% ACN and extracted peptides were dried under vacuum for 90 min at 45°C. For MALDI-TOF/TOF/MS analysis dried peptides were dissolved in peptide resuspension solution (0.1% TFA in 5% ACN) and desalted/concentrated using C18 zip tips (Merck Millipore) as per the manufacturer’s protocol. An equivalent volume (1 μL) of C18 zip tip purified peptides were mixed with a matrix solution (containing 10 mg/mL of α-cyano-4-hydroxy cinnamic acid matrix in 1 mL of 60% methanol-0.1% formic acid) and spotted on an Anchorchip target plate. Calibration was performed with TOF-Mix™ (LaserBio Labs, France) as an external standard. The peptide mass was analysed with a MALDI-TOF mass spectrometer (AXIMA Performance, SHIMADZU BIOTECH) in positive reflector ion mode and analysed by Shimadzu launch pad-MADLI MS software. Mono isotopic peak list (m/z range of 700-4000 kDa with S/N ratio over 10) was generated by MALDI MS software. Peptide mass fingerprints (PMFs) were analysed using online MASCOT server. When searching MASCOT, Swissprot database was mined against PMFs and MALDI-TOF/TOF/MS as instrument, *C. elegans* as the organism source, selected variable modification were carbamidomethylation of cysteine, oxidation, N-terminal acetylation and phosphorylation (S, T, Y) for methionine, where as a maximum of one missed cleavage sites was allowed and mass tolerance of 100 ppm per peptide was set as fixed modifications (Ananthi *et al*., 2011; Sethupathy *et al*., 2016).

#### 4.3.5 Bioinformatics analysis

Regulatory proteins identified from high throughput analysis were further subjected to bioinformatics analysis such as gene ontology (GO) classification in UniProt KB tool and interaction among them was assessed using STRING with medium confidence score 0.400. The gene enrichment score and functional annotation was generated using DAVID tool (Kamaladevi *et al*., 2017).

#### 4.3.6 *In vitro* embryo development toxicity assay

*C. elegans* embryos were obtained from gravid adult worms per protocol (Bianchi *et al*., 2006). Briefly, the worms from NGM agar plates were washed by Milli-Q water pellet down by centrifugation at 170 × g for 3 min. The pellets were lysed using the mixture of bleach and NaOH (Fresh Chlorox 5 mL, 10 N NaOH 1.25 mL and sterile H_2_O 18.75 mL) incubated for 5 min by giving gentle vortex several times and centrifuged at 170 × g for 3 min. The pellets and eggs were washed with egg buffer (118 mM NaCl, 48 mM KCl, 2 mM CaCl_2_, 2 mM MgCl_2_, and 25 mM Hepes pH 7.3) and centrifuged at 170 × g for 3 min. The pelleted eggs were resuspended and lysed in 2 mL of sterile egg buffer and 2 mL of a sterile 60% sucrose (in egg buffer) by vortexing. The suspension was centrifuged at 170 × g for 5-6 min. The eggs were collected from the top of the solution; isolated oocytes/eggs were treated in a 24-well microtitre plate containing *E. coli* OP50 (control) or isolated toxin proteins of *S.* Typhi (treated).

#### 4.3.7 Detection of oxidants and antioxidants

The L4 stage *C. elegans* treated to *E. coli* OP50 proteome, *S.* Typhi proteome and *E. coli* OP50 (bacteria), were washed several times thoroughly. Subsequently, the bacteria free worms were homogenized and protein concentration was maintained at 100 μg. Reactive oxygen species (ROS) was measured per Scherz□Shouval *et al*. (2007) to study the level of ROS in host during *S.* Typhi proteome exposure. The H_2_O_2_ level in cell lysate supernatant was measured as described earlier Wolff *et al*. (1994). SOD activity of *C. elegans* cell lysate supernatant (100 μg of protein) was measured as per Paoletti *et al*., 1990. Catalase activity of *C. elegans* was measured as described earlier Aebi *et al.*, 1984. *C. elegans* carbonyl content was measured as per Levine *et al*., 1990. Lipid peroxidation was determined as described earlier Ohkawa *et al*., 1979. Each experiment was performed in biological triplicates and the error bars represent the mean ± SD (\**p* < 0.05).

#### 4.3.8 ATP assay

Total intracellular ATP of *C. elegans* control and treated samples was measured as described earlier Chen *et al*., 1994. The protein concentration of the supernatant was determined and protein concentration was maintained at 500 μg (Bradford’s method) for all the three time points in triplicate. The ATP concentration in each sample was calculated according to the given formula:

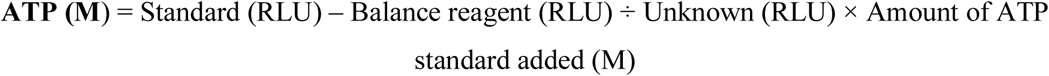

#### 4.3.9 Behavioural assay

Behavioural assay was performed to determine the impact of food source (*E. coli* OP50) to *S.* Typhi toxin proteome treated nematodes. The experiment was analysed *in vivo* using *C. elegans*. The age synchronized L4 stage N2 worms treated with toxin proteins (concentration 500 μg/mL, 1 mg/mL and 1.5 mg/mL) respectively, were incubated at 20°C for 12 hrs. After incubation the treated worms were washed with M9 buffer several times to remove traces of protein from the body and transferred to NGM agar plates seeded with food source *E. coli* OP50. The plates were incubated at 20°C and *C. elegans* life span and other physiological changes were monitored during the assays. The plates were scored for viability of *C. elegans* after every 2 hrs. Worms treated with *E. coli* OP50 were considered as control. Each experiment was performed in biological triplicates and the error bars represent the mean ± SD (\**p* < 0.05).

#### 4.3.10 Western Blotting

The protein content of whole cell extracts (at different time points of exposure) were prepared using lysis solution (containing 7 M urea, 2 M thiourea, 4% w/v CHAPS and 30 mM Tris-HCl, pH 8.5 along with protease inhibitor cocktail). Protein concentration was determined by Bradford assay. 100 μg of protein for each sample, was boiled in 5X LaemmLi buffer (0.3125 M Tris-HCl pH 6.8, 10% SDS, 50% glycerol, 0.005% bromophenol blue, 25% beta-mercaptoethanol) for 5 min followed by short spin at 1100 x g. The protein samples were subjected to SDS-PAGE followed by transfer on nitrocellulose membranes and subsequently, treated to the specific antibodies as described earlier (Durai *et al*., 2014). The antibodies used in this study were purchased from Santa Cruz Biotechnology, JNK-1[sc-571], p38 [sc-17852], HSF-1[sc-9144], HSF-90 [sc-1055], SGK-1 [sc-33774] and beta-actin purified mouse immunoglobulin (A1978) purchased form (Sigma-Aldrich) working concentration was kept at 1:1000 – 1:2000.

#### 4.3.11 Statistical analysis

All experiments were performed independently in triplicate. The statistical significance of data was analysed by one-way ANOVA and Duncan’s test (SPSS Chicago, IL, USA) at a significance level of *p* < 0.05.

## Abbreviations used in this resource article

2D-GE: Two Dimensional Differential Gel Electrophoresis,
DAF-21: abnormal Dauer Formation (protein),
JNK: C-Jun N-terminal Kinase,
MAPK: Mitogen Activated Protein Kinase,
MALDI-TOF: Matrix assisted laser desorption ionization-Time of Flight,
STRING: Search Tool for the Retrieval of Interacting

## Acknowledgement

We thank the Caenorhabditis Genetics Center, which is funded by the National Institute of Health, National Centre for Research Resources for providing the nematode strains. Mr. Dilawar Ahmad Mir gratefully acknowledges the University Grants Commission (UGC) of India for the financial assistance in the form of UGC-PF (F. No. 42-222/2013 (SR)). The computational facility provided by the Bioinformatics Infrastructure Facility, Alagappa University funded by the Department of Biotechnology, Ministry of Science and Technology, Government of India (Grant No.BT/BI/25/015/2012 (BIF)) are thankfully acknowledged. We are also grateful for the Instrumentation Facility provided by the Department of Science and Technology (DST), Government of India through DST PURSE [Grant No. SR/S9Z-415 23/2010/42(G)], DST FIST [Grant No. SR-FST/LSI-087/2008], UGC through SAP-DRS1 [Grant No. F. 3-28/2011(SAP-II)] and RUSA 2.0.

## Author contributions

DAM designed and performed the experiments and analysed the data. KB wrote the manuscript in consultation with DAM.

## Conflict of interest

The authors declare that they have no conflicts of interest with the contents of this article

## Manuscript Supplemental Figures

**Figure S1.** Proteomic analysis of *S. enterica* Typhi (MTCC 733). The *S. enterica* Typhi 500 mL culture was grown 12 hrs at 37°C, after incubation culture was centrifuged and collected *S. enterica* Typhi bacterial pellets was sonicated. The total cellular proteins of *S. enterica* Typhi were precipitated by adding the solid ammonium sulphate 20% – 80 %. Lane (1– 7) is ∼100 μg of *S.* Typhi precipitated protein from fraction (20% – 80%), respectively.

**Figure S2.** *C. elegans* total proteome (SDS-PAGE) after interaction against intact *S.* Typhi toxin proteins. Lane (1&3) is N2 control sample exposed with *E. coli* OP50 and Lane (2&4) is N2 sample treated with *S.* Typhi toxin proteins.

**Figure S3.** 2D gel electrophoreses images of *C. elegans* proteome. **A**).The upper panel representing the proteome of control nematodes fed on *E. coli* OP50.**B**).The lower panel representing the *C. elegans* total proteins treated by 1.5 mg/mL 50% *S.* Typhi protein fraction. The size of IP strip is 18 cm and the pI gradient is from 3 – 10. The experiment was performed in triplicates.

## Manuscript Supplemental tables

**Table S1.** List of downregulated proteins present in *C. elegans* (control sample) identified using MALDI-TOF/TOF/MS.

**Table S2**. List of upregulated proteins present in *C. elegans* (treated sample) identified using MALDI-TOF/TOF/MS

## References

Aballay, A., Drenkard, E., Hilbun, L.R., and Ausubel, F.M. (2003). Curr. Biol. 13, 47–52.

Aebi, H. (1984). Catalase *in vitro*. Methods Enzymol. 105,121–126.

Ananthi, S., Santhosh, R.S., Nila, M.V., Prajna, N.V., Lalitha, P., and Dharmalingam, K. (2011). Experimental eye research, 92, 454–463.

Balasubramanian, V., Sellegounder, D., Suman, K. and Krishnaswamy, B., (2016). Journal of proteomics, 145, pp.141–152.

Bianchi, L. and Driscoll, M., (2006).[WormBook: The Online Review of C. elegans Biology (Internet)].

Boscher, C., Dennis, J.W., and Nabi, I.R. (2011Curr. Opin. Cell Biol. 23, 383–392.

Bradford, M.M. (1976). Anal. Biochem. 72, 248–254.

Braverman, N.E., and Moser, A.B. (2012).. Biochim. Biophys. Acta, Mol. Basis Dis. 1822, 1442–1452.

Cezairliyan, B., Vinayavekhin, N., Grenfell-Lee, D., Yuen, G.J., Saghatelian, A., and Ausubel, F.M. (2013). PLoS pathog. 9, p.e1003101

Chen, F. and Cushion, M.T., (1994). Journal of clinical microbiology, 32(11), pp.2791–2800

Chen, W.W., Yi, Y.H., Chien, C.H., Hsiung, K.C., Ma, T.H., Lin, Y.C., Lo, S.J., and Chang, T.C. (2016). Sci. Rep. 6, 32021.

Durai, S., Pandian, S.K., and Balamurugan, K. (2011). J. Basic Microbiol. 51, 243–252.

Durai, S., Singh, N., Kundu, S., and Balamurugan, K. (2014). Proteomics. 14,1820–1832.

Esterbauer, H., and Cheeseman, K.H. (1990). Methods Enzymol. 186, 407–421.

Espinosa-Diez, C., Miguel, V., Mennerich, D., Kietzmann, T., Sánchez-Pérez, P., Cadenas, S., and Lamas, S. (2015). Redox biol. 6, 183–197.

Etheridge, T., Oczypok, E.A., Lehmann, S., Fields, B.D., Shephard, F., Jacobson, L.A., and Szewczyk, N.J. (2012). PLoS genet. 8, e1002471.

Fouad, A.D., Pu, S.H., Teng, S., Mark, J.R., Fu, M., Zhang, K., Huang, J., Raizen, D.M., and Fang-Yen, C. (2017). G3.7, 1811–1818.

Frydman, J. (2001. Annu. Rev. Biochem. 70, 603-647.

Galán, J.E. (2001). Annu. Rev. Cell Dev. Biol. 17, 53–86.

Gal-Mor, O., Boyle, E.C., and Grassl, G.A. (2014). Front. Microbiol. 5, 391.

Green, R.A., Kao, H.L., Audhya, A., Arur, S., Mayers, J.R., Fridolfsson, H.N., Schulman, M., Schloissnig, S., Niessen, S., Laband, K., and Wang, S. (2011). Cell. 145, 470–482.

Griffitts, J.S., Huffman, D.L., Whitacre, J.L., Barrows, B.D., Marroquin, L.D., Müller, R., Brown, J.R., Hennet, T., Esko, J.D., and Aroian, R.V. (2003). J. Biol. Chem. 278, 45594–45602.

Hellerer, T., Axäng, C., Brackmann, C., Hillertz, P., Pilon, M., and Enejder, A. (2007). PNAS. 104,14658–14663.

Hodgkin, J., Kuwabara, P.E., and Corneliussen, B. (2000). Curr. Biol. 10,1615–1618.

Huffman, D.L., Bischof, L.J., Griffitts, J.S., and Aroian, R.V. (2004). Int. J. Med. Microbiol. 293, 599–607.

Hunt, P.R. (2017). J. Appl. Toxicol. 37, 50–59.

JebaMercy, G., Prithika, U., Lavanya, N., Sekar, C., and Balamurugan, K. (2015). Gene. 558, 159–172.

JebaMercy, G., Durai, S., Prithika, U., Marudhupandiyan, S., Dasauni, P., Kundu, S. and Balamurugan, K. (2016). Journal of proteomics, 145, pp.81–90.

Kamaladevi, A., and Balamurugan, K. (2016). RSC Adv. 6, 30070–30080.

Kamaladevi, A., and Balamurugan, K. (2017). Cell. Infect.Microbiol. 7, 393.

Kasai, K.I., and Hirabayashi, J. (1996). J. Biochem. 119, 1–8.

Kesika, P., Karutha Pandian, S., and Balamurugan, K. (2011). Scand. J. Infect. Dis. 43, 286–295.

Kesika, P., and Balamurugan, K. (2012). Biochim. Biophys. Acta, Proteins Proteomics. 1824, 1449–1456.

Kho, M.F., Bellier, A., Balasubramani, V., Hu, Y., Hsu, W., Nielsen-LeRoux, C., McGillivray, S.M., Nizet, V., and Aroian, R.V. (2011). PLoS One, 6, e29122.

Levine, R.L. (1990). Meth. Enzymol. 186, 465–478.

Li, H., Ren, C., Shi, J., Hang, X., Zhang, F., Gao, Y., Wu, Y., Xu, L., Chen, C., and Zhang, C. (2010). Proteome Sci. 8, 49.

Liu, Y., Zhang, Q., Hu, M., Yu, K., Fu, J., Zhou, F., and Liu, X. (2015). Infect. Immun. 83, 2897–2906.

Leung, M.C., Williams, P.L., Benedetto, A., Au, C., Helmcke, K.J., Aschner, M., and Meyer, J.N. (2008). Toxicol. Sci. 106, 5–28.

Lindblom, T.H., and Dodd, A.K. (2006). J. Exp. Zool. A. Ecol. Genet. Physiol. 305, 720–730.

Lyczak, J.B., Cannon, C.L., and Pier, G.B. (2000). Microb. Infect. 2, 1051–1060.

Marsh, E.K., and May, R.C. (2012). Appl. Environ. Microbial.78, 2075–2081.

Mohri-Shiomi, A., and Garsin, D.A. (2008). J. Biol. Chem. 283, 194–201.

Ohkawa, H., Ohishi, N., and Yagi, K. (1979). Anal. Biochem. 95, 351–358.

Otey, C.A., Rachlin, A., Moza, M., Arneman, D. and Carpen, O., (2005) International review of cytology, 246, pp.31–58.

Paoletti, F., and Mocali, A. (1990). Meth. Enzymol. 186, 209–220.

Park, G.E., Oh, H.N. and Ahn, S.Y., (2009). Bulletin of the Korean Chemical Society, 30(9), pp.2032–2038.

Park, J.W., Lee, S.G., Song, J.Y., Joo, J.S., Chung, M.J., Kim, S.C., Youn, H.S., Kang, H.L., Baik, S.C., Lee, W.K., and Cho, M.J. (2008). Electrophoresis, 29, 2891–2903.

Petriv, I., Pilgrim, D.B., Rachubinski, R.A., and Titorenko, V.I. (2002). Physiol. Genomics. 10, 79–91.

Popham, J.D., and Webster, J.M. (1979). Caenorhabditis elegans. Environ. Res. 20, 183–191.

Powers, M.V., Jones, K., Barillari, C., Westwood, I., Montfort, R.L.V., and Workman, P. (2010). Cell Cycle, 9, 1542–1550.

Prithika, U., Deepa, V., and Balamurugan, K. (2016). Innate Immun. 22, 466–478.

Pryor, W.A. (1989). Free Radic. Biol. Med. 7, 177–178.

Ray, A., Rentas, C., Caldwell, G.A., and Caldwell, K.A. (2015). Neurosci. Lett. 584, 23–27.

Rogalski, T.M., Gilbert, M.M., Devenport, D., Norman, K.R., and Moerman, D.G. (2003). Genetics. 163, 905–915.

Schmutz, C., Ahrné, E., Kasper, C.A., Tschon, T., Sorg, I., Dreier, R.F., Schmidt, A., and Arrieumerlou, C., (2013). Mol. Cell. Proteomics. 12, 2952–2968.

Schlesinger D (ed.) (1975) D. American Society for Microbiology, Washington, D.C.

Scherz□Shouval, R., Shvets, E., Fass, E., Shorer, H., Gil, L., and Elazar, Z. (2007). EMBO J. 26, 1749–1760.

Schouest, K., Zitova, A., Spillane, C., and Papkovsky, D.B. (2009). Environ. Toxicol. Chem. 28, 791–799.

Sethupathy, S., Prasath, K.G., Ananthi, S., Mahalingam, S., Balan, S.Y., and Pandian, S.K. (2016). J. Proteomics. 145, 112–126.

Shi, X., Tarazona, P., Brock, T.J., Feussner, I., and Watts, J.L. (2016). J.Lipid Res. 57, 265–275.

Sivamaruthi, B.S., and Balamurugan, K. (2014). Ind. J. Microbial. 54, 52–58.

Smith, H. and Pearce, J.H., (1972). Br. J. Pharmacol. 146, 769–780.

Soti, C., Nagy, E., Giricz, Z., Vígh, L., Csermely, P. and Ferdinandy, P., (2005). British journal of pharmacology, 146(6), pp.769-780.

Stiernagle Theresa (2006): Maintenance of Caenorhabditis elegans WormBook: The Online Review of Caenorhabditis elegans Biology.

Vigneshkumar, B., Pandian, S.K., and Balamurugan, K. (2012). Arch. Microbiol. 194, 229–242.

Wählby, C., Conery, A.L., Bray, M.A., Kamentsky, L., Larkins-Ford, J., Sokolnicki, K.L., Veneskey, M., Michaels, K., Carpenter, A.E., and O’Rourke, E.J. (2014). Methods. 68, 492–499

Wang, B., Wang, H., Xiong, J., Zhou, Q., Wu, H., Xia, L., Li, L., and Yu, Z. (2017). Sci. Rep.7, 14170.

Wingfield, P. (2001). Curr. Protoc. Protein Sci. A-3F.

Wolff, S.P. (1994). Meth. Enzymol. 233, 182–189.

Yang, R.Y., Rabinovich, G.A., and Liu, F.T. (2008). Expert Rev. Mol. Med. 10, e17.

